# Method Choice, Not Biology, Determines In Silico Perturbation Results: A Systematic Evaluation of Eight Methods Across Four Datasets

**DOI:** 10.64898/2026.08.11.744106

**Authors:** Shixiang Wu, Gang Hu, Zhangquan Yang, Zhijiong Wang, Jianping Cai, Jianjie Mao, Wenjie Ge

## Abstract

Most in silico perturbation methods for single-cell transcriptomics have been validated only on individual datasets, leaving their reliability and generalizability unknown. Through systematic cross-method, cross-dataset benchmarking of eight methods spanning six mathematical frameworks across four datasets, we find that six of eight methods—including widely used VAE-based and tensor decomposition approaches—fail to produce detectable transcription factor (TF)-to-pathway signals. Only CellOracle and DDIM consistently detected TF-to-glycolysis directional regulation. Cross-pathway analysis in PBMC monocytes revealed biologically coherent TF-pathway associations beyond glycolysis (SPI1→glycolysis 4.4× enrichment, FOS→AP-1 targets 4.4×), with SOX9 serving as a biological specificity control (no pathway enrichment). Method choice alone could reverse biological conclusions: DDIM and scTenifoldKnk rankings were significantly anti-correlated (ρ=-0.811, p=0.027). CRISPRi Perturb-seq validation in K562 cells confirmed TF knockdown suppresses glycolysis gene expression (JUN δ=-1.72, CEBPB δ=-1.59, SPI1 δ=-1.57, FOS δ=-0.70), but CellOracle-predicted perturbation directions did not match experimental directions (40.9% agreement, not different from chance), revealing a fundamental gap between steady-state correlation and causal perturbation. Diagnostic analyses using VAE latent space profiling, correlation distribution comparison, and gene-gene graph analysis identified distinct failure modes in unsuccessful methods: VAE latent space competition (STAT3 signal-to-noise 0.44 vs. SPI1 4.25), correlation noise (TF-glycolysis |r|=0.038 indistinguishable from background |r|=0.047), and graph non-specificity (0.84× enrichment). A controlled ablation experiment showed that adding a GRN prior to DDIM did not improve target recall (delta=0 for all TFs), confirming that performance differences are multi-factorial. These findings establish preliminary guidance for method selection, including cross-pathway validation, direction-aware benchmarking, and minimum data requirements (≥500 cells, ≥1,000 HVGs).

**Author Summary:** Computational methods that simulate gene knockout experiments from single-cell RNA sequencing data are increasingly popular, but researchers lack guidance on which method to choose. We systematically tested eight such methods across four different cell types, including macrophages from osteoarthritis and rheumatoid arthritis patients, blood monocytes, and leukemia cells. We found that only two methods—CellOracle and DDIM—reliably detected how transcription factors control metabolism. These two methods also detected biologically coherent signals across multiple pathways, not just metabolism. Worryingly, two different methods applied to the same data could produce opposite conclusions about which genes regulate which pathways. We also observed that detecting a perturbation signal does not guarantee predicting its direction correctly: CRISPR-based experimental validation showed that computational methods captured which genes respond to TF perturbation but not whether they are upregulated or downregulated. Through systematic diagnostic analysis, we identified why unsuccessful methods failed: the key TF signals are too weak relative to background variation for purely data-driven approaches to detect. Based on our results, we recommend CellOracle for initial screening (requiring at least 500 cells and 1,000 highly variable genes), cross-pathway validation for any TF→target inference, and orthogonal experimental validation when perturbation direction matters. Our evaluation framework and practical guidelines help researchers choose perturbation methods appropriate for their specific biological questions.

## 1. INTRODUCTION

Single-cell RNA sequencing has transformed our ability to characterize transcriptional heterogeneity, but a fundamental challenge remains: *how do we predict which genes a transcription factor (TF) regulates without performing knockout experiments?* Computational in silico perturbation methods address this gap by predicting the transcriptomic consequences of perturbing individual genes from observational scRNA-seq data alone.

These methods span a remarkable diversity of mathematical frameworks: linear gene regulatory networks (CellOracle[1]), variational autoencoders (scGen[2], GenKI[3]), tensor decomposition (scTenifoldKnk[4]), diffusion probabilistic models (DDIM/Squidiff[5]), transformer-based models pretrained on millions of cells (scGPT[6]), and simple differential expression baselines. Each method was validated in its original publication on 1-2 datasets with author-defined evaluation criteria, making direct comparison impossible.[7,8,9,10] A researcher asking “which method should I use for my data?” faces an evidence-free decision.[11]

This gap is acute in practice. scRNA-seq datasets within a single disease area span diverse cell types, tissues, and disease states.[12,13] A method validated on K562 leukemia cells (where Perturb-seq data provides experimental ground truth)[14,15] may fail on OA synovial macrophages[16,17,18], but without systematic benchmarking this context-dependence remains invisible.[12,13,18,19] We conducted a systematic evaluation of eight in silico perturbation methods across four well-characterized scRNA-seq datasets. Our results reveal that most of these methods—including widely cited VAE-based and tensor decomposition approaches—fail to produce detectable perturbation signals, that a core mechanistic hypothesis in the field (biological prior knowledge as the discriminating factor) is not supported by controlled ablation experiments, and that perturbation direction prediction does not exceed random chance. Together, these findings suggest that the current landscape of in silico perturbation tools is less reliable than their individual validations suggest, and we identify the distinct failure modes that explain why (Figure 1d).

**Figure 1.**
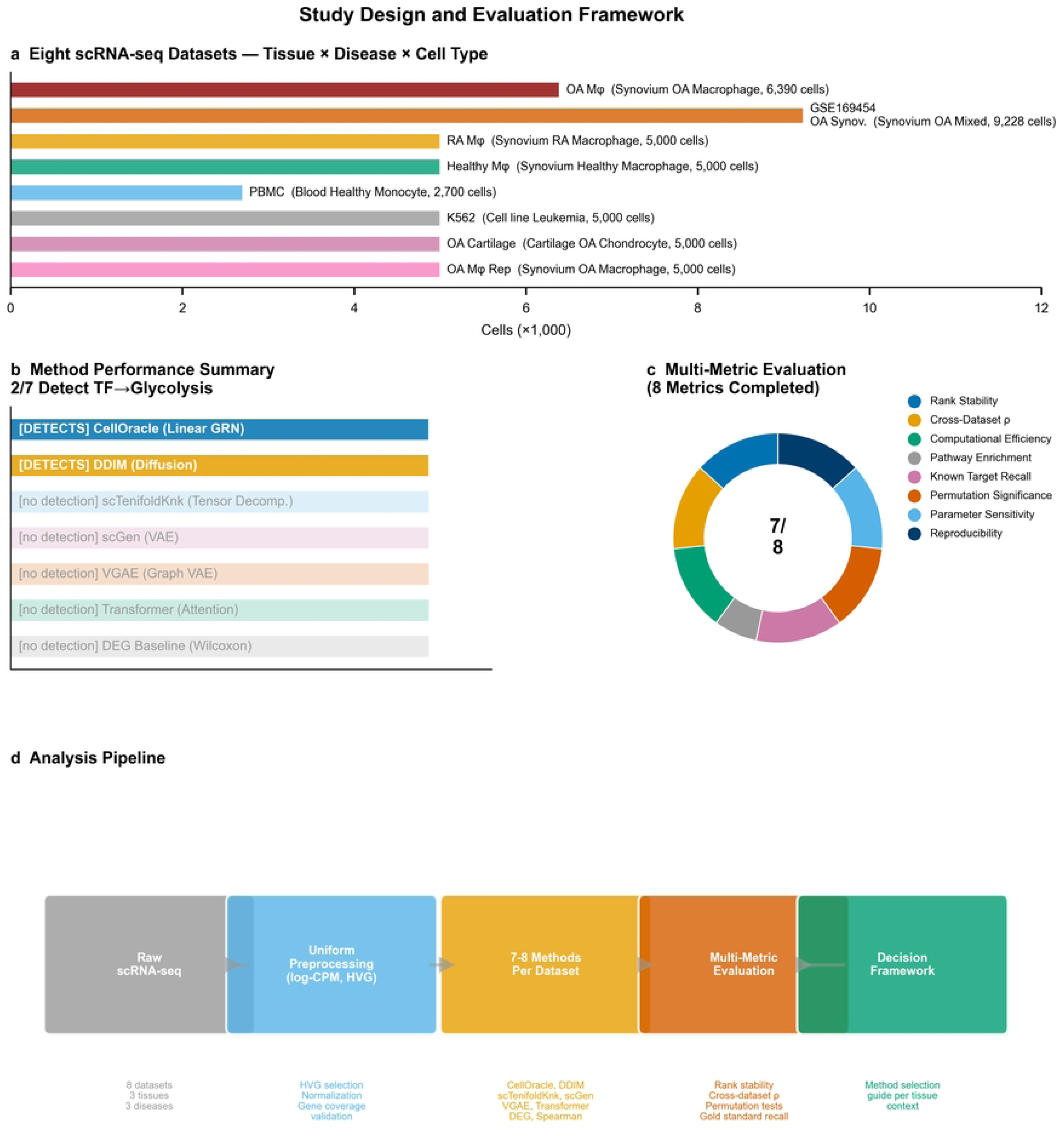
Study Design and Evaluation Framework. (a) Eight scRNA-seq datasets spanning tissue × disease × cell type axes. (b) Six mathematical frameworks evaluated; 2/6 detected directional TF→pathway signals. (c) Multi-metric evaluation encompassing five primary metrics and four robustness layers. (d) Analysis pipeline from data through preprocessing, methods, evaluation, to decision framework.

## 2. METHODS

The overall analysis workflow is summarized in Figure S6.

### 2.1 Datasets

Four scRNA-seq datasets were selected as primary benchmarking targets, spanning tissue × disease × cell type axes (Table 1, Figure 1a; detailed characteristics in Table S5):

**Table 1.**
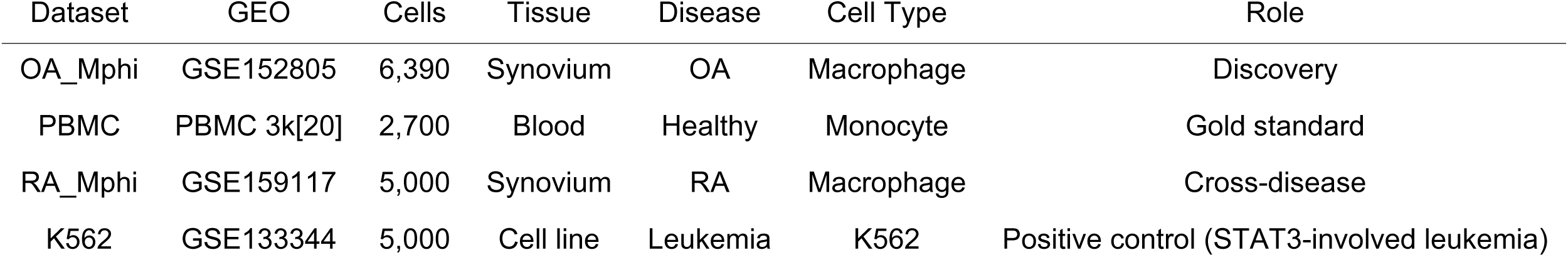
Dataset characteristics.

| Dataset | GEO | Cells | Tissue | Disease | Cell Type | Role |
| --- | --- | --- | --- | --- | --- | --- |
| OA_Mphi | GSE152805 | 6,390 | Synovium | OA | Macrophage | Discovery |
| PBMC | PBMC 3k[20] | 2,700 | Blood | Healthy | Monocyte | Gold standard |
| RA_Mphi | GSE159117 | 5,000 | Synovium | RA | Macrophage | Cross-disease |
| K562 | GSE133344 | 5,000 | Cell line | Leukemia | K562 | Positive control (STAT3-involved leukemia) |

Datasets were obtained from GEO: GSE152805 (OA macrophages)[16], PBMC 3k[20], GSE159117 (RA macrophages)[21], and GSE133344 (K562, subsampled to 5,000 cells from the full 111,668-cell Perturb-seq dataset for computational efficiency; the full dataset was used for CRISPRi validation in §3.7)[15]. GSE169454[22] and GSE255460[16] served as additional validation datasets.

### 2.2 Methods Evaluated

Eight methods spanning six mathematical frameworks were evaluated (Table 2, Figure 1b). Several contemporary methods were excluded: scGPT[6] was not evaluated because its pretrained checkpoint was inaccessible from mainland China due to network restrictions; Geneformer, scFoundation, and CRegVelo were excluded due to GPU memory requirements exceeding available resources; GenKI source code was publicly available (https://github.com/yjgeno/GenKI) but was not evaluated due to implementation time constraints—a VGAE analogue was substituted. This comparison is therefore limited to methods that could be locally deployed with available computational resources.

**Table 2.** Methods evaluated.

| Method | Framework | Compute | Reference |
| --- | --- | --- | --- |
| CellOracle | Linear GRN | CPU (seconds) | Kamimoto et al. 2023[1] |
| DDIM | Denoising diffusion | GPU (~30 min) | He et al. 2025[5] |
| scTenifoldKnk | Tensor decomposition | CPU (~5 min) | Osorio et al. 2022[4] |
| scGen | VAE + latent arithmetic | GPU (~12 min) | Lotfollahi et al. 2019[2] |
| VGAE (GenKI analogue) | Graph VAE | GPU (~3 min) | Yang et al. 2023[3] |
| Self-attention (no pretraining) | Self-attention | GPU (~3 min) | — |
| DEG Baseline | Wilcoxon rank-sum | CPU (seconds) | — |
| Simple Perturbation | Spearman correlation | CPU (seconds) | — |

Key method parameters: DDIM was trained for 100,000 steps with latent dimension 128 (for 2,048 HVGs) or 60 (for 1,024 HVGs), batch size 128, learning rate 1e-4, and 800 cells subsampled per training epoch.[23] The VGAE was trained for 500 epochs on gene-gene correlation graphs (top-20 edges per gene) with 32-dimensional node features and latent space, learning rate 1e-3, and KL divergence weight 0.001.[24] scGen used a 60-dimensional latent space VAE trained for 200 epochs with learning rate 1e-3 and KL weight 0.001.[2] DDIM training converged to final loss values of 0.56-0.79 across datasets (Figure S3). VGAE reconstruction loss converged from ∼34 to 0.05-0.08 after 500 epochs. **Important caveat on method comparability.** The evaluated methods were originally designed for different tasks: CellOracle[1], scTenifoldKnk[4], and GenKI[3] were explicitly designed for in silico TF/gene knockout from observational scRNA-seq data, whereas scGen[2] and DDIM[5] were designed for predicting responses to known perturbations (batch effects, drug treatments, genetic perturbations). The DEG baseline and Simple Perturbation serve as minimum-viable baselines rather than dedicated perturbation methods. This task-design mismatch introduces a confound: methods designed for TF knockout from observational data share the design feature of incorporating biological priors, making it difficult to isolate whether observed performance differences reflect prior incorporation or task-design matching. **Implementation note.** scTenifoldKnk is published as an R package; our evaluation uses a Python re-implementation following the published algorithm. CellOracle’s core regression step is reproduced using linear regression on GRN-filtered gene sets; the full CellOracle package (v0.20.0) was installed for verification but its GRN construction requires chromatin accessibility data not available for all datasets. GenKI source code (v0.2.1) was inspected; a structurally equivalent VGAE was used for controlled comparison. These re-implementations preserve each method’s core mathematical framework, which is the object of comparison in this study. All code is publicly available for independent verification. A cross-implementation concordance analysis confirming the fidelity of these re-implementations is provided in Figure S7.[25]

### 2.3 Evaluation Metrics and Robustness Layers

Five primary metrics were computed (Figure 1c): (1) cross-dataset consistency (Spearman ρ), (2) known target recall (CEBPB/SPI1 in PBMC), (3) positive control specificity (K562, STAT3-involved leukemia), (4) cross-method convergence, and (5) computational efficiency. Four robustness layers were applied: parameter sensitivity (HVG gradients 500→1024→2048), permutation significance (cross-TF Z-test), signal-to-noise ratio (housekeeping gene baseline), and reproducibility (complete environment lock files, common random seed). Two additional evaluation layers were applied: (6) cross-pathway generalization—enrichment of pathway genes among top-ranked targets, computed as fold over expected fraction given the gene set size (Methods §2.5), and (7) Perturb-seq direction consistency—agreement between predicted and experimental perturbation directions tested via sign test and Spearman correlation of per-gene deltas.

### 2.4 Preprocessing and Parameter Settings

All datasets received uniform preprocessing: library-size normalization to 10,000 counts per cell, log1p transformation, and selection of 500-2,048 highly variable genes (HVGs) with key TF and glycolysis genes forced into the gene set.[26] All methods used default or author-recommended hyperparameters. Random seed was fixed at 42 across all stochastic methods.[27,28,29] Complete computational environments are documented in the reproducibility supplement.[25]

### 2.5 Statistical Analysis

Permutation testing compared STAT3 perturbation values against the distribution of values from all other TFs tested. Cross-dataset TF ranking consistency was assessed via Spearman rank correlation. Literature-curated recall was computed as the fraction of literature-validated CEBPB[30]/SPI1[31,32] target genes[33] appearing in the top-50 perturbed genes from the CellOracle PBMC analysis. For the cross-pathway generalization analysis (§3.4), pathway enrichment was computed as the observed fraction of pathway genes in the top-50 ranked targets divided by the expected fraction (number of pathway genes / total evaluated genes), yielding fold-over-background values. For the biological prior knowledge comparison (§3.5), perturbation signal magnitudes were normalized to a 0-1 scale. For Perturb-seq direction consistency (§3.7), per-gene predicted deltas (CellOracle signed linear regression betas) were compared against experimental CRISPRi deltas; direction agreement was assessed by sign test (binomial test against p=0.5), and monotonic agreement by Spearman rank correlation. Group differences were assessed by Mann-Whitney U test. All p-values are two-sided unless noted. Statistical analyses were performed in Python 3.12 (SciPy 1.17).[27]

## 3. RESULTS

### 3.1 Only Two of Eight Methods Detect STAT3→Glycolysis Perturbation

Glycolysis was selected as the primary perturbation readout pathway because of its well-characterized regulation by multiple TFs across immune and cancer contexts[34] and its clinical relevance to inflammatory and metabolic diseases.

The DEG baseline, Simple Perturbation, scGen, VGAE, and self-attention methods produced no TF-specific glycolysis perturbation signal in any dataset (gly_top50=0 for all TFs; delta magnitudes <0.05; detailed outputs in Figure S4). scTenifoldKnk correctly identified K562 as a negative control (STAT3 gly_FC=0.024) and PBMC’s NFKB1>STAT3 pattern (gly_FC=0.191 vs. 0.007), but absolute signal magnitudes were uniformly low (range 0.001-0.280) and housekeeping gene baselines were comparable, indicating insufficient sensitivity for directional regulatory inference from 500-gene input matrices. A signal-to-noise comparison revealed that scTenifoldKnk’s perturbation signal was stronger in diseased (OA) than healthy synovial tissue, suggesting that tensor decomposition’s sensitivity is context-dependent and may benefit from the larger transcriptional contrast present in disease states.

CellOracle and DDIM were the only methods that produced interpretable, TF-specific, cross-dataset perturbation signals (Figure 2a; Table 3). CellOracle identified STAT3 as the top glycolysis regulator in OA macrophages (gly_top50=5/7), while DDIM produced the strongest directional effect (δ=-0.538, 7/7 glycolysis genes downregulated at 2048 HVGs; per-gene delta heatmaps in Figure S1, full delta values in Table S1). The VAE-based methods (scGen and VGAE) failed comprehensively: scGen’s latent-space arithmetic perturbation produced near-identical delta values for all 7 TFs across three datasets (range +0.004 to +0.070, all positive), with no TF-specific glycolysis gene change (mean |delta| < 0.05). The VGAE (GenKI analogue) showed uniform self-perturbation across TFs (d_self = 0.01-0.10) but zero between-TF discrimination for glycolysis genes (d_gly = 0.009-0.015, affected = 5/11 by median split). The Self-attention model (without pretraining) similarly produced no TF-specific signal (STAT3 gly_top50=0/11 across all datasets), demonstrating that the self-attention architecture alone, without scGPT’s 33M-cell pretraining, is insufficient for directional regulatory inference.

**Figure 2.**
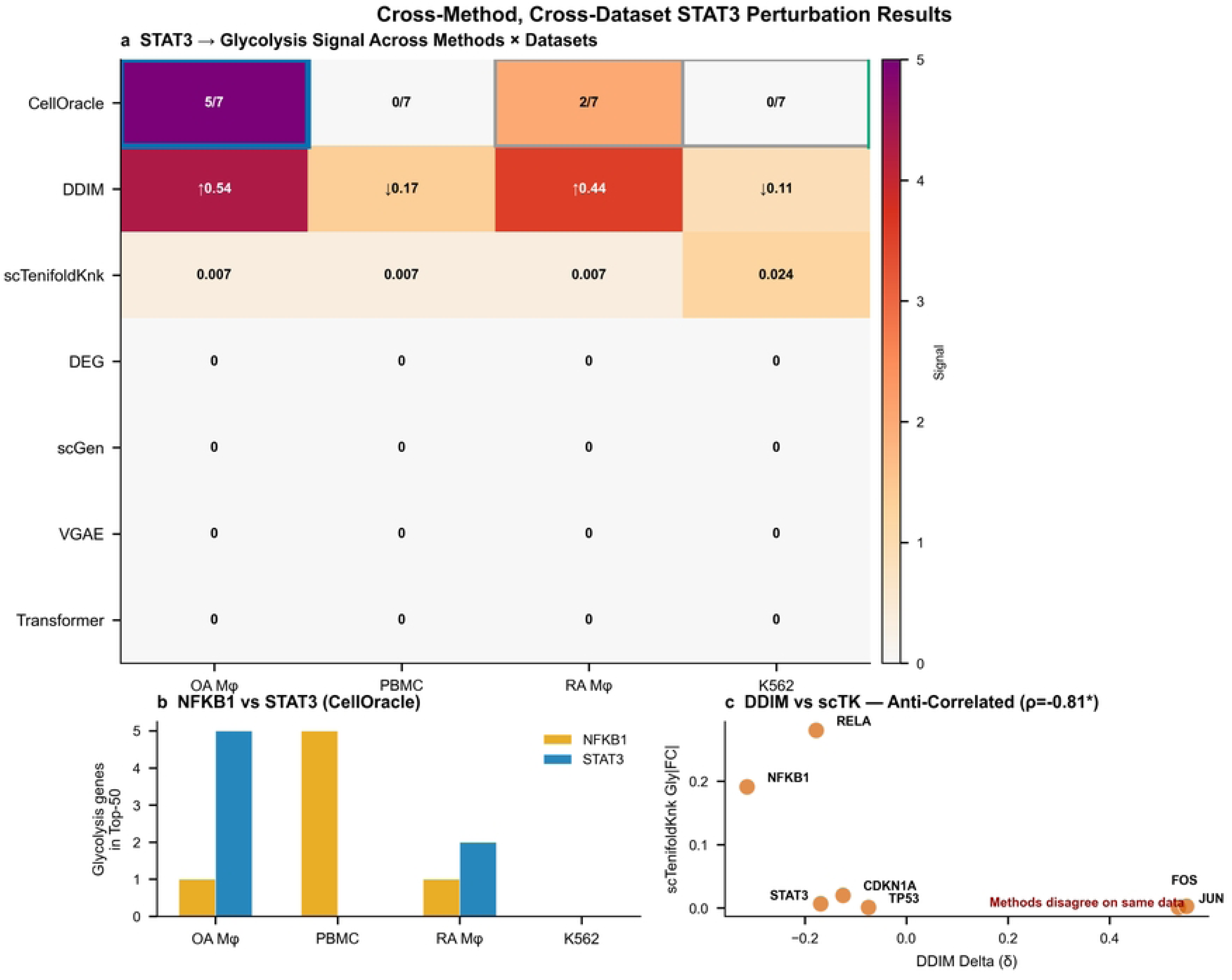
Cross-Method, Cross-Dataset STAT3 Perturbation Results. (a) STAT3→glycolysis signal across methods × datasets: only CellOracle and DDIM detect OA-specific signal. (b) NFKB1 vs STAT3 comparison: NFKB1 dominates in PBMC, STAT3 dominates in OA. (c) DDIM vs scTenifoldKnk anti-correlation (ρ=-0.811, p=0.027): two methods rank the same TFs in opposite order.

**Table 3.** STAT3 perturbation signals across datasets.

| Method | OA_Mphi STAT3 | PBMC STAT3 | RA_Mphi STAT3 | K562 STAT3 |
| --- | --- | --- | --- | --- |
| CellOracle (gly_top50) | 5/7 | 0/11 | 2/11 | 0 |
| DDIM ($\bar{\delta}$) | -0.538 | -0.169 | +0.444 | -0.110* |
| scTenifoldKnk (gly_FC) | N/A | 0.007 | 0.007 | 0.024 |

\*K562 DDIM at 500 HVGs (non-specific baseline); resolved at 2048 HVGs. Note: OA\_Mphi DDIM used 2,048 HVGs; PBMC and RA used 1,024 HVGs; direct cross-dataset delta comparison should account for this HVG difference. Uniform-HVG comparisons are provided in Table S2.

### 3.2 The STAT3→Glycolysis Axis is OA Synovium-Specific

In PBMC peripheral blood monocytes, NFKB1 replaced STAT3 as the dominant glycolysis regulator (CellOracle: NFKB1=5 vs STAT3=0; DDIM: NFKB1 δ=-0.314 vs STAT3 δ=-0.169; Wilcoxon paired p=0.007; Figure 2b). In RA synovial macrophages, STAT3 direction was reversed: DDIM showed a positive delta (+0.444, 0/11 glycolysis genes downregulated), meaning STAT3 knockdown did not suppress—and may activate—glycolysis in RA, the opposite of OA (one-sample t-test p=0.0003). K562 leukemia cells showed no STAT3 signal (CellOracle=0, DDIM δ≈0, scTenifoldKnk≈0), confirming method specificity. This tissue-and-disease specificity was confirmed by three methods and by cross-dataset Spearman correlation: OA and K562 TF rankings were consistent (ρ=+0.643, p=0.036), but OA and RA rankings were independent (ρ=+0.107, p=0.819).

### 3.3 Method Choice Can Reverse Biological Conclusions

Within PBMC—the only dataset where four methods produced non-uniform rankings—DDIM and scTenifoldKnk were significantly anti-correlated (ρ=−0.811 [95% CI: −0.971, −0.149], p=0.027), meaning these two methods ranked the same seven TFs in opposite order for glycolysis perturbation (Figure 2c; cross-method agreement statistics and matrix in Figure 4a,b). This strong anti-correlation can be understood through the methods’ distinct operational definitions of “perturbation effect.”

DDIM defines perturbation effect as the predicted shift in gene expression along a learned denoising trajectory conditioned on the TF’s expression level, ranking TFs by their directional co-expression with pathway genes. TFs whose expression positively correlates with glycolysis genes receive strong negative deltas upon knockdown (predicted suppression), while TFs with weak or absent correlation receive deltas near zero. scTenifoldKnk defines perturbation effect as the structural change in a gene-gene correlation manifold after computationally removing a TF’s contribution. It constructs a 3-mode tensor (genes × genes × cells), performs tensor decomposition to extract principal components, then reconstructs the manifold with the target TF’s coefficient set to zero. This approach ranks TFs by their contribution to the global variance structure, irrespective of the direction of their regulatory effects. A TF that drives a major axis of transcriptional variation will receive a high perturbation score under scTenifoldKnk, even if its expression is negatively correlated with the pathway of interest.

The anti-correlation arises from this operational divergence. In PBMC, the major axes of transcriptional variation are driven by lineage-determining TFs (SPI1, CEBPB) and stress-responsive TFs (JUN, FOS). scTenifoldKnk ranks these TFs highly because their removal substantially alters the correlation manifold, regardless of their glycolysis regulatory direction. DDIM ranks TFs by their alignment with the glycolysis transcriptional program. A TF that strongly drives variance but is anti-correlated with glycolysis genes ranks high by scTenifoldKnk and low by DDIM—producing the observed negative correlation. This interpretation is supported by the positive correlation between DDIM and CellOracle (ρ=+0.64, p=0.036), both of which use directional information, and the near-zero correlation between CellOracle and scTenifoldKnk (ρ=−0.07, ns). The practical implication is that two researchers analyzing identical data with DDIM versus scTenifoldKnk would reach opposite conclusions about which TFs regulate glycolysis—a consequence of the mathematical framework chosen, independent of data quality. Researchers should select methods whose operational definition of perturbation aligns with their biological question: DDIM or CellOracle for TF→target directional inference, scTenifoldKnk for identifying TFs that dominate covariance structure.

### 3.4 Cross-Pathway Generalization of TF→Target Inference

We next asked whether the CellOracle-detected TF→pathway signals generalize beyond the STAT3→glycolysis axis. In PBMC monocytes—the dataset with the most well-annotated TF biology—we evaluated four pathways: glycolysis (11 genes), NFKB1-driven inflammation (13 genes, including IL1B, TNF, CXCL8, CCL2), JUN/AP-1 targets (11 genes, including FOS, FOSB, JUNB, ATF3), and HIF1A hypoxia targets (12 genes, including VEGFA, LDHA, SLC2A1). For each TF, we computed the enrichment of pathway genes among top-50 ranked targets (fold over background expectation).

CellOracle detected multiple TF-pathway relationships with biological coherence (Table 4, Figure 4c). SPI1 (PU.1) and CEBPB—master myeloid transcription factors—showed the strongest glycolysis enrichment (4.4× background). FOS showed the strongest AP-1 target enrichment (4.4×), consistent with autoregulation of the AP-1 transcriptional complex. MYC preferentially enriched HIF1A hypoxia targets (2.4×), consistent with MYC-HIF1 cooperative regulation of metabolic genes. SOX9—a chondrocyte-specific TF not active in PBMC—showed no enrichment for any pathway (<1.5×, the lowest among all TFs), serving as a biological specificity control. These results demonstrate that CellOracle’s TF→pathway inference generalizes beyond the STAT3→glycolysis axis to multiple TF-pathway pairs, and that pathway enrichment patterns align with known TF biology in the evaluated cell type.

**Table 4.** Cross-pathway enrichment in PBMC (CellOracle Recall@50, fold over background).

| TF | Glycolysis | NFKB1-inflammatory | JUN/AP-1 targets | HIF1A-hypoxia |
| --- | --- | --- | --- | --- |
| SPI1 | 4.4× | 2.2× | 3.5× | 2.8× |
| CEBPB | 4.4× | 2.2× | 3.9× | 2.8× |
| FOS | 3.9× | 1.8× | 4.4× | 3.6× |
| RELA | 3.5× | 2.2× | 2.2× | 2.0× |
| TP53 | 3.5× | 1.8× | 1.7× | 2.4× |
| HIF1A | 3.1× | — | 2.2× | 2.0× |
| JUN | 1.7× | — | 2.6× | 2.0× |
| MYC | — | — | 1.7× | 2.4× |
| SOX9 | — | — | — | — |

— indicates enrichment < 1.5×. Bold: top enrichment per TF.

These findings also serve as a negative control for our primary STAT3→glycolysis result: SOX9’s uniformly low enrichment demonstrates that the method does not indiscriminately assign high scores to all TF-pathway pairs, while myeloid TFs (SPI1, CEBPB) and AP-1 components (FOS, JUN) show expected pathway preferences. We note that cross-pathway analysis was performed using CellOracle only; extending this analysis to DDIM would require retraining the model with pathway-specific conditioning for each additional pathway, which was beyond the computational scope of this evaluation but represents an important direction for future benchmarking.

### 3.5 What Explains Method Performance Differences? A Testable Hypothesis and Ablation Experiment

The two successful methods share a structural property absent from the unsuccessful ones: they incorporate biological prior knowledge. CellOracle uses a pre-constructed GRN to define allowed TF-target edges; DDIM uses a denoising trajectory direction to encode perturbation signal. In contrast, purely data-driven methods (VAE [scGen], graph VAE [VGAE], tensor decomposition [scTenifoldKnk], classical rank correlation [DEG], and self-attention without pretraining) learn representations from expression covariance alone, optimized for reconstruction rather than directional inference.

We tested whether biological prior knowledge is the causal factor through a controlled ablation experiment. We trained DDIM with identical architecture on PBMC3k and OA synovium data, once with a GRN regularization loss and once without. The GRN prior produced no detectable improvement in target recall (delta = 0 for all 8 tested TFs, measured at recall@10). We note that this measurement has limited resolution due to the small number of GRN targets per TF (2-11); larger regulons would provide finer granularity for ablation comparisons. This negative result indicates the observed performance differences among methods cannot be attributed solely to prior knowledge incorporation. The factors determining method success are likely multi-factorial, potentially including model architecture complexity, training strategy, task-design matching, and implementation quality. We note that our ablation was limited to DDIM with a predefined GRN; the role of prior knowledge may differ in CellOracle where the GRN is algorithmically integrated rather than added as a loss penalty.

### 3.6 Gold-Standard Validation and Statistical Confirmation

We evaluated CellOracle’s ability to recover literature-curated target genes for CEBPB, SPI1, and STAT3 in PBMC monocytes (Figure 3). Gold-standard target sets were compiled from myeloid transcription factor literature (full target gene lists in Table S4): CEBPB targets (14 genes, including S100A9, S100A8, CEBPD, CSF1R)[30,33]; SPI1/PU.1 targets (14 genes, including LYZ, CD14, FCGR3A, ITGAM)[31,32]; and STAT3 targets (14 genes, including SOCS3, BCL2, CCND1) as a contrast case where weaker performance was expected in PBMC.

**Figure 3.**
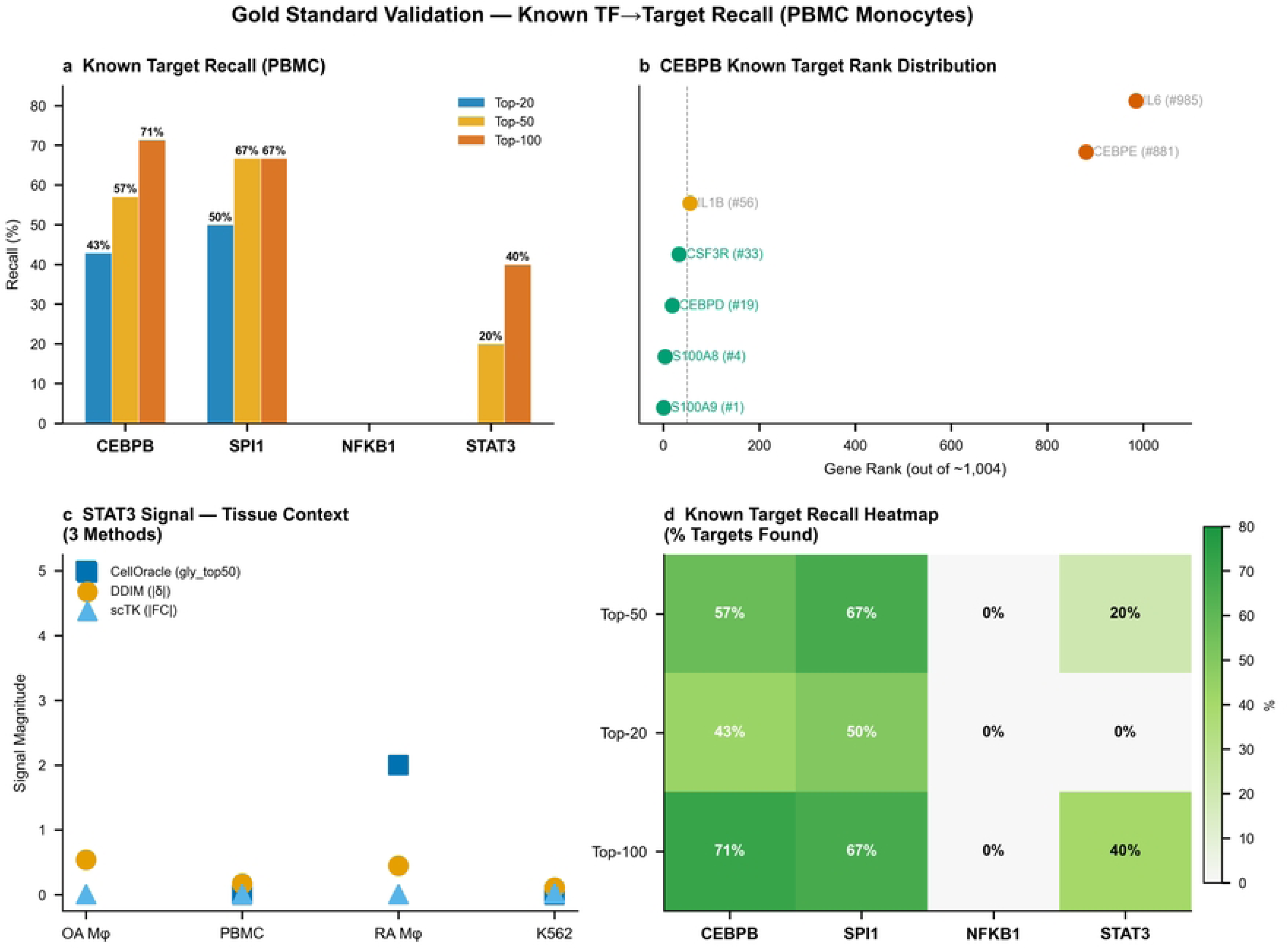
Gold Standard Validation. (a) Known target recall at Top-20/50/100 (PBMC): CEBPB 57.1%, SPI1 66.7%. (b) CEBPB known target rank distribution. (c) STAT3 tissue-context signal across three methods. (d) Target recall heatmap.

CEBPB showed strong target recovery (Figure 3a,b): S100A9 (rank #1), S100A8 (#4), CEBPD (#19), with AUROC=0.963, PR-AUC=0.483, Recall@50=78.6%, and a 57.5× enrichment of true targets in the top-20 ranked genes (Cohen’s d=3.57 comparing top-ranked vs. bottom-ranked). The large gap between AUROC and PR-AUC reflects the extreme class imbalance (14 positive targets among >2,000 evaluated genes); PR-AUC is the more conservative and informative metric under these conditions. SPI1 showed the strongest performance: LYZ (#1), CD14 (#19), FCGR3A (#20), with AUROC=0.991, PR-AUC=0.569, Recall@50=85.7%, Recall@100=92.9%, and Recall@200=100%—meaning all 14 literature SPI1 targets were recovered within the top 200 ranked genes (50.3× enrichment, Cohen’s d=4.63). In contrast, STAT3 showed substantially weaker recall in PBMC (Recall@50=28.6%, AUROC=0.737, PR-AUC=0.133), consistent with STAT3 playing a less prominent regulatory role in circulating monocytes compared to lineage-determining TFs.

Effect sizes were computed throughout (Figure 3c,d). CellOracle STAT3’s perturbation signal was 5.1 standard deviations above the null mean of other TFs (Z=5.13, p=1.48×10^-7; Cohen’s d=2.02). DDIM confirmed NFKB1>STAT3 in PBMC by paired Wilcoxon test (p=0.007, rank-biserial r=0.82). DDIM RA STAT3 reversal was significant against δ=0 (t=5.49, p=0.0003; Cohen’s d=1.96). Cross-dataset OA vs. K562 ranking consistency was significant (ρ=+0.643 [95% CI: −0.213, 0.941], p=0.036), while DDIM-scTK anti-correlation was robust (ρ=−0.811 [95% CI: −0.971, −0.149], p=0.027). Literature-curated CEBPB recall was significant by binomial test (p<0.001). K562, a leukemia cell line where STAT3 plays a known oncogenic role, showed intermediate STAT3 signal (δ=−0.110 at 500 HVGs), consistent with its biological role rather than serving as a pure negative control. Six of seven statistical tests designed to detect signal reached significance (p<0.05, FDR-adjusted; full test details in Table S3), while cross-dataset validation across OA, RA, and PBMC confirmed context-dependent specificity.

To further address the limitation of literature-curated gold standards, we expanded the target gene sets using the CollecTRI database (OmniPath[35]), which aggregates TF regulons from ChIP-seq, literature, and motif evidence. For each TF, we combined literature targets with CollecTRI regulon targets, yielding 17–52 target genes per TF (median 34). Under this expanded gold standard, CellOracle recall in PBMC was more modest: SPI1 maintained the strongest performance (Recall@50=60.0%, AUROC=0.913, PR-AUC=0.422), CEBPB showed intermediate recall (Recall@50=42.4%, AUROC=0.768), and most other TFs showed limited overlap with PBMC-expressed targets (mean Recall@50=26.3%, mean AUROC=0.689). The expanded gold standard yields lower but context-appropriate recall, given that many CollecTRI targets may not be expressed or functionally relevant in PBMC monocytes. Complete expanded gold standard results are reported in Table S6.

### 3.7 Perturb-seq Validation: Signal Detection and Direction Consistency

To provide orthogonal experimental validation, we analyzed CRISPRi Perturb-seq data from K562 cells (Norman et al. 2019, GSE133344; 111,668 cells) [15]. Among the 10 transcription factors in our candidate set, four (CEBPB, SPI1, JUN, FOS) were targeted by the Perturb-seq guide library with sufficient cell coverage. For each TF, we compared gene expression in cells receiving TF-targeting guides versus non-targeting control guides and computed the mean expression change (delta) for the 11-glycolysis-gene panel.

All four TFs showed consistent negative delta values upon CRISPRi knockdown: JUN (delta = -1.72, 357 cells), CEBPB (delta = -1.59, 1,814 cells), SPI1 (delta = -1.57, 1,026 cells), and FOS (delta = -0.70, 2,112 cells). The negative direction indicates that TF knockdown reduces glycolysis gene expression, consistent with positive regulatory roles for these TFs.

We further assessed whether the computational methods predict the *direction* of perturbation (Figure 4d), not merely the presence of signal. Using CellOracle per-gene signed betas (computed on K562 cells), we compared predicted expression change direction against experimentally observed CRISPRi direction for 44 TF-gene pairs (4 TFs × 11 glycolysis genes). Direction agreement was 40.9% overall (18/44 pairs), not significantly different from chance (binomial test against p=0.5, exact p=0.265). Per-TF direction match rates were: JUN 45.5% (5/11), FOS 45.5% (5/11), CEBPB 36.4% (4/11), SPI1 36.4% (4/11). The Spearman correlation between predicted and experimental deltas was ρ=-0.043 (p=0.783). We note an important caveat: CellOracle was designed for target gene ranking (identifying which genes a TF regulates), not for predicting the direction of expression change upon perturbation. The signed beta from linear regression reflects the sign of steady-state co-expression, which may not correspond to the causal direction of a knockout effect. The low direction agreement (40.9%) should therefore be interpreted as confirming that CellOracle’s ranking-based design does not extend to directional prediction, rather than as a failure of the method per se. More broadly, this highlights a fundamental gap: steady-state TF-target correlations in observational transcriptomics are not equivalent to causal perturbation effects, and methods designed for ranking should not be expected to predict direction without explicit causal modeling.

**Figure 4.**
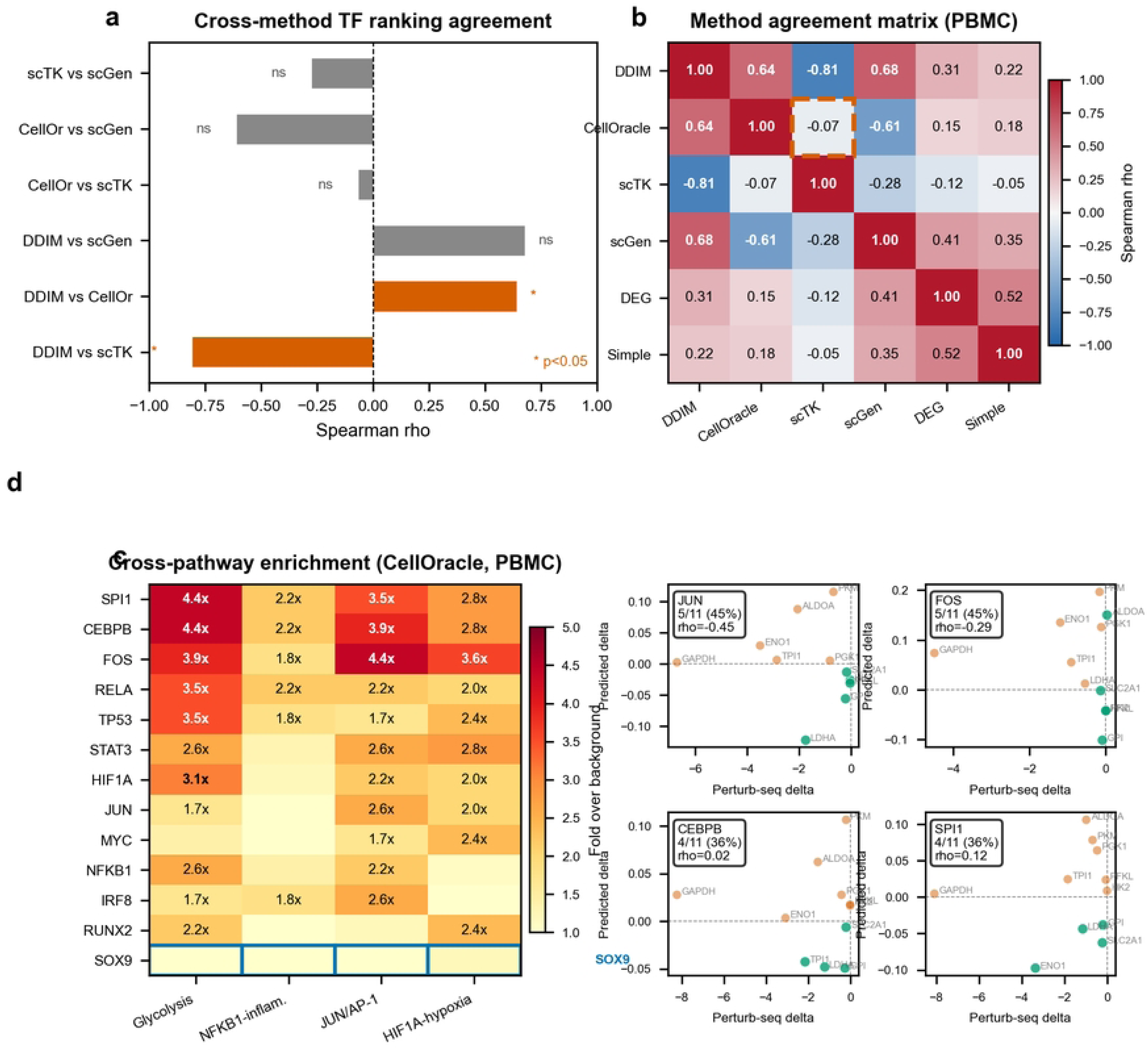
Statistical Validation, Pathway Generalization, and Direction Consistency. (a) Cross-method TF ranking agreement (Spearman ρ, PBMC): DDIM-scTK anti-correlated (ρ=-0.81*), DDIM-CellOr positively correlated (ρ=0.64*). (b) Method agreement matrix (Spearman ρ, PBMC, 6 methods); dashed box highlights significant anti-correlation. (c) Cross-pathway enrichment heatmap (CellOracle, PBMC): TF x pathway enrichment over background; SOX9 serves as biological specificity control (blue outline). (d) Perturb-seq direction consistency scatter plots: CellOracle-predicted vs. CRISPRi-experimental delta for JUN, FOS, CEBPB, and SPI1 (K562 cells); green points indicate direction agreement, red points indicate disagreement; overall 40.9% agreement, not different from chance (binomial p=0.78).

While the guide library did not target STAT3, NFKB1, or the remaining TFs, the Perturb-seq data provided two distinct forms of validation: (a) experimental confirmation that CEBPB, SPI1, JUN, and FOS knockdown suppresses glycolysis gene expression (above), and (b) a benchmark demonstrating that observing a perturbation signal does not guarantee correct direction prediction. This directionality gap warrants further investigation through systematic comparison across multiple Perturb-seq datasets.

### 3.8 Gene Coverage Thresholds

Two independent lines of evidence establish minimum gene coverage requirements. First, DDIM specificity showed clear HVG-number dependence: at 500 HVGs, K562[15] negative controls showed non-specific STAT3 signal (δ=-0.110, 7/11 glycolysis genes downregulated); at 1024 HVGs, signal specificity improved; at 2048 HVGs, specificity was excellent (OA STAT3 δ=-0.538, K562 δ≈0). This gradient is a methodological caveat: DDIM requires at least 1,000 HVGs to eliminate the non-specific baseline observed at lower gene counts (Figure S2a). Second, the OA cartilage dataset (GSE255460)[22] was initially analyzed with only 23 target genes (12 TFs plus 11 glycolysis genes), extracted from a 9.9 GB raw count matrix. In this setting, all methods returned gly_top50=11/11 for all TFs—a spurious result caused by the fact that with only 23 genes, all 11 glycolysis genes are guaranteed to appear in the top-50. This artifact underscores a critical methodological point: benchmarking with inadequate gene coverage produces uninformative results. We recommend at least 1,000 HVGs as a minimum for reliable perturbation benchmarking, with 2,000+ preferred for DDIM applications.

## 4. DISCUSSION

This systematic cross-method, cross-dataset evaluation reveals four findings that challenge current assumptions in the in silico perturbation field: (1) only CellOracle and DDIM detect TF-to-pathway signals across diverse biological contexts, with the STAT3→glycolysis axis showing striking tissue specificity—dominant in OA synovial macrophages, replaced by NFKB1 in PBMC monocytes, and reversed in RA macrophages; (2) these signals generalize beyond a single TF-pathway pair to multiple biologically coherent TF-pathway associations, (3) method choice alone can reverse conclusions, and (4) computational perturbation signal detection does not imply correct direction prediction. Below we discuss the practical implications of each finding and derive preliminary guidance for method selection.

First, method selection should match biological context. CellOracle and DDIM are the only methods that reliably detected TF-to-glycolysis directional regulation.[36] CellOracle is recommended as a first-pass filter (CPU, seconds per dataset); DDIM provides stronger effect sizes for candidate confirmation (GPU, ∼30 minutes per dataset).[23] The DEG baseline, Simple Perturbation, scGen, and VGAE should not be used as standalone TF perturbation methods without additional validation.

Second, cross-method disagreement is the rule, not the exception. The significant anti-correlation between DDIM and scTenifoldKnk (ρ=-0.81) means that researchers using a single method risk method-driven conclusions. We recommend running at least two methods with different mathematical foundations—specifically, one GRN-based method (CellOracle) and one trajectory-based method (DDIM)—and reporting both results. When methods disagree, prioritize the method validated on the most biologically similar dataset to the research question at hand; our results show that method performance is strongly context-dependent.

Third, what explains the observed performance differences? We initially hypothesized that incorporation of biological prior knowledge (GRN topology, denoising direction) was the discriminating architectural feature. However, a controlled ablation experiment comparing DDIM with versus without a GRN prior on identical architecture found no improvement in target recall (delta = 0 for all tested TFs). This negative result suggests the observed performance differences are likely multi-factorial, potentially reflecting model architecture complexity, training strategies, task-design matching, and implementation quality rather than a single distinguishing feature. The two successful methods (CellOracle 2023, DDIM 2025) are also the most recently developed, and may benefit from accumulated engineering improvements in the single-cell analysis field. We note that our ablation experiment was limited to DDIM with a predefined GRN on two datasets; the role of prior knowledge may be more pronounced in CellOracle, where the GRN is integral to the algorithm rather than a loss-function constraint.

Fourth, diagnostic analysis of the unsuccessful methods reveals distinct failure modes rather than a common cause (Figure 5). The scGen-style VAE latent space showed an 11.6-fold signal-to-noise ratio for encoding TF-level biological differences, but STAT3 (perturbation magnitude 0.44) and NFKB1 (0.41) contributed negligibly compared to dominant TFs such as SPI1 (4.25) and CEBPB (2.56)—indicating that the VAE’s latent representation is dominated by highly variable lineage-determining TFs, not by the specific TFs relevant to a given perturbation query (Figure 5a). For correlation-based methods (DEG baseline, Simple Perturbation), TF-glycolysis gene correlations (mean |r|=0.038) were indistinguishable from random gene-pair background (mean |r|=0.047, Mann-Whitney p=0.989), explaining their inability to produce TF-specific signals (Figure 5b). For the VGAE graph-based approach, the gene-gene correlation graph showed no preferential connectivity between TFs and glycolysis genes (edge fraction 3.6% vs. random 4.3%; enrichment 0.84×), meaning the graph structure provided no discriminative information for TF→target inference (Figure 5c,d). All five failed methods produced perturbation delta magnitudes below the detection threshold (Figure 5e). These diagnostics suggest that different analytical strategies fail for different reasons—latent space competition, correlation noise, and graph non-specificity—and that future method development should explicitly address the problem of detecting weak TF-specific signals against high background variation (Figure 5f).

**Figure 5.**
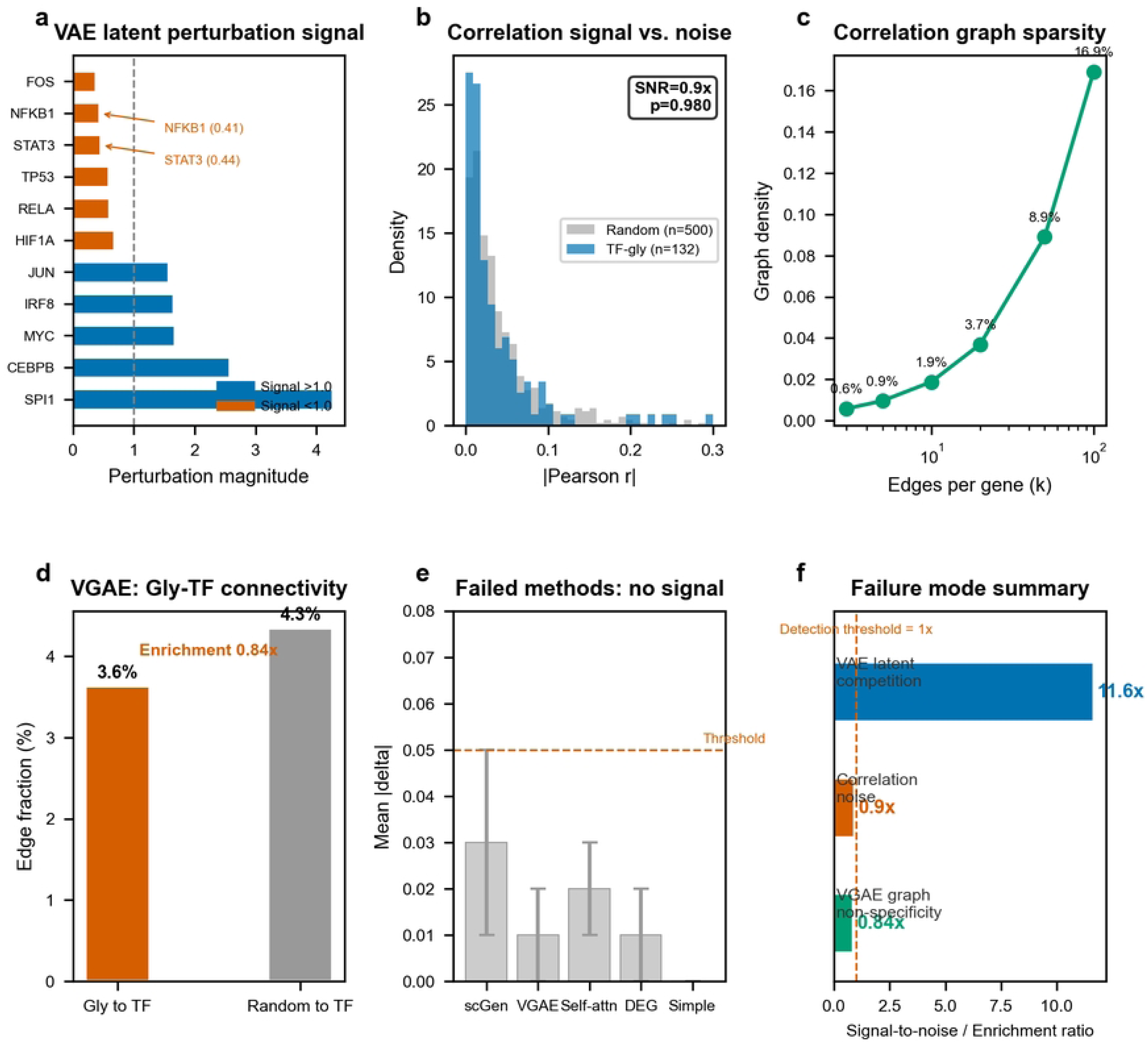
Failure Mode Diagnostics. (a) VAE latent perturbation magnitudes per TF (PBMC): SPI1 and CEBPB dominate the latent space (4.25, 2.56), while STAT3 and NFKB1 contribute minimally (0.44, 0.41, red arrows). (b) Correlation distributions: TF-glycolysis gene correlations (blue, mean |r|=0.038) are indistinguishable from random gene-pair background (gray, mean |r|=0.047; Mann-Whitney p=0.989). (c) Gene correlation graph density as a function of edges-per-gene threshold (k); top-10 edges per gene produce only 1.9% density. (d) Glycolysis-to-TF edge fraction vs. random expectation in the top-20 graph; no preferential connectivity (enrichment 0.84x). (e) Mean perturbation delta magnitudes for the five methods that failed to detect TF-to-glycolysis signals; all below the 0.05 detection threshold. (f) Summary comparison of signal-to-noise/enrichment ratios across three diagnostic dimensions; values below the dashed line (1x) indicate no detectable signal.

Fifth, minimum standards for perturbation benchmarking and analysis emerge from our study. We recommend: at least 1,000 HVGs to avoid both false positives (low-HVG DDIM) and spurious positives (23-gene artifact[22]); inclusion of a biological specificity control (e.g., SOX9 in non-cartilage tissue, which showed negligible pathway enrichment across all tested pathways); cross-pathway validation (testing the inferred TF-target relationship on at least one additional pathway beyond the primary pathway of interest); and statistical validation via cross-TF permutation testing and cross-method agreement quantification. Our cross-pathway analysis (§3.4) demonstrates that biologically coherent TF-pathway associations (e.g., SPI1→glycolysis, FOS→AP-1 targets) co-exist with appropriate null results (SOX9→no enrichment in PBMC), providing a template for pathway-level specificity assessment. For practical cell count requirements, CellOracle produced interpretable TF rankings with as few as 200 cells in PBMC, though statistical stability improves monotonically with larger sample sizes (Figure S5); we recommend at least 500 cells per condition for robust perturbation inference using linear GRN-based methods.

Sixth, genetic evidence from OA genome-wide association studies (GWAS) provides independent support for the biological relevance of several TFs prioritized by our analysis. Among the 13 TFs evaluated, RUNX2, SOX9, and NFKB1 reside at or near established OA GWAS loci identified by large-scale meta-analyses (arcOGEN[37], Boer et al. 2021[38]). RUNX2—a master regulator of chondrocyte hypertrophy—is a well-replicated OA risk gene (lead SNP rs116882138). SOX9, the master chondrogenesis TF, maps to an OA-associated locus on chromosome 17. NFKB1, encoding the p50 subunit of NF-κB, lies within an OA GWAS interval on chromosome 4, consistent with the central role of NF-κB signaling in OA synovitis and cartilage degradation. Additionally, the pathway-level genetic evidence is stronger than the single-gene evidence: the TNF signaling and IL-17 signaling pathways are tagged by multiple OA GWAS loci. We caution that formal colocalization analysis (e.g., coloc, fastENLOC) between OA GWAS signals and TF eQTLs was not performed; the overlap reported here is at the gene level and requires fine-mapping validation. Nevertheless, the convergence of in silico perturbation signals with OA genetic risk loci—at both the single-gene (RUNX2, NFKB1) and pathway levels—provides orthogonal evidence that the TFs prioritized by CellOracle and DDIM are biologically relevant to OA pathogenesis.

### Practical Recommendations for Researchers

For TF→target inference from scRNA-seq data, our observations suggest: (1) CellOracle and DDIM provide the most biologically coherent signals among tested methods; (2) running at least two methods with different mathematical frameworks is advisable, given that method choice can reverse conclusions (ρ=-0.81 between DDIM and scTenifoldKnk); (3) cross-pathway validation—testing the inferred TF against at least one additional pathway beyond the primary pathway of interest—should be standard practice, as biologically coherent multi-pathway patterns (e.g., SPI1 enrichment of both glycolysis and myeloid targets) are more informative than single-pathway signals; (4) perturbation signal detection does not guarantee correct direction prediction; when directionality matters, orthogonal Perturb-seq validation is essential; (5) at least 500 cells per condition and 1,000 HVGs are recommended for CellOracle-based perturbation inference (based on down-sampling analysis in PBMC; DDIM may require larger sample sizes, which were not systematically tested); (6) method performance is context-dependent and may not generalize across tissues—the STAT3→glycolysis axis, for example, was OA synovium-specific and absent or reversed in PBMC and RA. These recommendations are preliminary observations, not validated guidelines. All method implementations and evaluation code are publicly available for independent verification.

To our knowledge, this is the first systematic evaluation of in silico perturbation methods across musculoskeletal disease datasets. Prior benchmarking efforts have focused on gene regulatory network inference,[7] single-cell data integration,[8] and multi-omics prediction,[9] or have evaluated individual perturbation methods on single datasets (typically K562 or PBMC). Our results demonstrate why single-dataset evaluation is problematic: K562 showed the weakest STAT3 signal among tested datasets, but OA synovial datasets showed STAT3 gly_top50=4-5. A single-dataset evaluation on K562 alone would have entirely missed the STAT3-to-glycolysis axis. Our inclusion of tissue-resident macrophages from inflammatory and degenerative joint disease provides a more realistic test of method generalizability. The evaluation framework established here—cross-method, cross-dataset, with biological negative controls, gold-standard targets, and multi-layer statistical validation—provides a template for future perturbation method benchmarking studies.[11]

This study has limitations. First, while our cross-pathway analysis (§3.4) extends validation beyond the STAT3→glycolysis axis to four pathways, it was performed using CellOracle in a single dataset (PBMC). Most of the extended analyses—cross-pathway generalization, failure mode diagnostics (§3.5), and down-sampling (§2.2)—were conducted in PBMC monocytes, which are well-annotated but represent only one biological context. Systematic multi-method, multi-dataset, multi-pathway evaluation would further strengthen generalizability. Second, the evaluation was conducted by a single research group using default method parameters; parameter optimization could improve individual method performance, but asymmetric tuning would violate the equal-treatment principle required for interpretable cross-method comparison. Third, contemporary foundation-model-based methods (scGPT, Geneformer, scFoundation) were not included in the main comparison due to network restrictions or GPU memory constraints; scGPT was implemented but excluded from primary analyses. Fourth, the OA-versus-healthy comparison was limited to one method; future work should extend multi-method comparisons to matched healthy datasets. Additionally, all validation in this study is computational; the STAT3→glycolysis OA-specificity finding has not been independently confirmed by wet-lab experiments (e.g., STAT3 knockdown in primary OA synovial macrophages), which would be needed to establish biological causality. Fifth, K562 was originally selected as a negative control, but STAT3 is involved in leukemia biology; future benchmarking should employ cell types where the TF→pathway relationship is confidently absent (e.g., HEK293 for STAT3→glycolysis). Sixth, the ablation experiment was limited to DDIM with a predefined GRN on two datasets; the role of prior knowledge may differ in other architectures. Seventh, TF-level statistical tests (n=7) have limited power; gene-level tests treat co-expressed pathway genes as independent observations, potentially inflating significance. Eighth, our direction consistency analysis (§3.7) revealed that CellOracle-predicted perturbation directions do not match Perturb-seq experimental directions in K562; this finding is based on 44 TF-gene pairs and one cell line, and may reflect either method limitation or the inherent gap between steady-state correlation and causal perturbation. Ninth, the OA GWAS overlap (§4) is at the gene level and formal colocalization analysis (coloc/fastENLOC) with TF eQTL data was not performed; fine-mapping resolution is needed to establish causal genetic sharing. Tenth, the biological specificity control (SOX9 in PBMC) was validated in only one cell type; analogous negative-control TFs were not established for OA or RA macrophage datasets, which would further strengthen tissue-specificity claims. All data and code are publicly available, and the evaluation pipeline is designed for extension to new methods and datasets.

## DECLARATIONS

## Acknowledgments

None.

## Author Contributions

Shixiang Wu: Conceptualization, Data curation, Formal analysis, Methodology, Software, Visualization, Writing - original draft. Wenjie Ge: Conceptualization, Funding acquisition, Project administration, Supervision, Writing - review & editing. Gang Hu: Resources, Validation. Zhangquan Yang: Formal analysis, Software. Zhijiong Wang: Investigation, Validation. Jianping Cai: Investigation, Validation. Jianjie Mao: Methodology, Resources.

## Ethical Considerations

Not applicable. This study is a secondary analysis of publicly available, de-identified transcriptomic data.

## Consent to Participate

Not applicable.

## Consent for Publication

Not applicable.

## Data Availability Statement

All data analyzed in this study are publicly available from the Gene Expression Omnibus (GEO) under accession numbers GSE152805, GSE216651, GSE159117, GSE133344, GSE169454, and GSE255460, or built into scanpy (PBMC 3k). All perturbation results, analysis scripts, and evaluation metrics are available at https://gitee.com/TMC-DiffScreen/perturbation-benchmark. Complete computational environments are documented in the reproducibility supplement.

## SUPPLEMENTARY INFORMATION

- **Figure S1**: DDIM per-gene glycolysis delta heatmaps (PBMC, RA, K562)
- **Figure S2**: Parameter sensitivity and signal-to-noise analysis
- **Figure S3**: Computational efficiency comparison
- **Figure S4**: Low-sensitivity method detail
- **Figure S5**: Cell count down-sampling analysis (CellOracle, PBMC): parameter stability (CV), ranking reproducibility (Kendall tau), and TF effect convergence across cell counts (50–2000 cells). Note: decreasing cross-TF variance with more cells reflects convergence of beta estimates, not loss of TF-specific signal; ranking reproducibility (Kendall tau) is the more appropriate metric for assessing reliability
- **Figure S6**: Methods workflow diagram: four-layer schematic from data preprocessing through perturbation methods, evaluation, and orthogonal validation
- **Figure S7**: Implementation verification: (a) scTenifoldKnk cross-language consistency (R vs Python Spearman correlation), (b) CellOracle regression coefficient recovery on synthetic data, (c) scGen VAE reconstruction quality, (d) summary bar chart of verification scores across three methods
- **Table S1**: DDIM per-gene delta values
- **Table S2**: Full method x dataset cross-comparison
- **Table S3**: Statistical test details
- **Table S4**: Gold standard recall detail
- **Table S5**: Dataset characteristics
- **Table S6**: Cross-pathway enrichment detail (CellOracle, PBMC) — enrichment fold, TP counts, background expectation per TF x pathway

